# Purified CBD and CBD-rich full-spectrum *Cannabis sativa* extract potentiate the angiogenic paracrine function of umbilical cord derived mesenchymal stem cells

**DOI:** 10.64898/2026.07.04.736503

**Authors:** Javiera Fontecilla-Escobar, Karina Flores-Montero, Hilde Harb Buzza, Rodrigo Acuña Astudillo, Ignacio Hernández, Ana I. Bellomo Peraza, Eleonora Elhalem, Gregorio Bigatti, Diego O. Croci, Marcelo Ezquer, Maria Celeste Ruete

## Abstract

**Background:** Chronic and non-healing wounds remain a major clinical challenge with limited therapeutic options. Angiogenesis and inflammation are central to tissue repair, and mesenchymal stem cells (MSC) contribute to these processes through their trophic and immunomodulatory secretome. Cannabidiol (CBD) exhibits antioxidant and immunomodulatory properties. However, whether CBD-rich *Cannabis sativa* extract stimulate MSC toward a pro-angiogenic secretome remains unclear.

**Purpose:** This study aims to determine whether purified CBD or a phytochemically CBD-rich full spectrum extract stimulate umbilical cord-derived human MSC (UC-hMSC) to secrete pro-angiogenic factors and enhance endothelial responses relevant to wound healing.

**Methods:** UC-hMSC were preconditioned with either purified CBD or a CBD-rich full-spectrum extract. Transcriptional changes were assessed by qPCR. The functional impact of the resulting secretomes was evaluated *in vitro* using HUVEC-based proliferation and tube formation assays, and *in vivo* through the chick chorioallantoic membrane assay. To explore underlying mechanisms, we examined HIF-1α stabilization and VEGFA release in UC-hMSC, and VEGFR-2/ERK signaling in HUVEC.

**Results:** Purified CBD and full-spectrum CBD extract preconditioned UC-hMSC secretomes, increased HUVEC proliferation, tube formation, and enhanced vascular branching in the CAM assay. Mechanistic analyses indicated activation of the HIF-1α/VEGF axis in UC-hMSC, and ERK1/2 activation in HUVEC that was sensitive to VEGFR-2 blockade.

**Conclusion:** Purified CBD and CBD-rich full-spectrum extract prime UC-hMSC toward a pro-angiogenic secretome that promotes endothelial activation and neovascularization. These findings suggest that cannabinoid-based preconditioning of UC-hMSC involves the HIF-1α/VEGF axis and VEGFR-2/ERK signaling pathways in endothelial cells, supporting further investigation of this approach in wound healing and regenerative therapies.

## Introduction

Chronic and non-healing wounds remain a major clinical challenge, with high morbidity, substantial healthcare costs, and limited therapeutic options (Olsson et al., 2019). The prevalence of these wounds is increasing, driven by aging populations and the rising incidence of comorbidities such as diabetes, obesity, and vascular disease. Wound healing is a complex, tightly regulated process involving hemostasis, inflammation, proliferation, and remodeling (Mamun et al., 2024; Okonkwo and DiPietro, 2017). Among these phases, angiogenesis during the proliferative stage is critical to restore tissue oxygenation and nutrient supply, and its dysregulation underlies both impaired wound closure and pathological scarring (Goldberg and Diegelmann, 2017; Korntner et al., 2019; Okonkwo and DiPietro, 2017; Veith et al., 2019). Accordingly, therapeutic strategies aimed at enhancing controlled angiogenesis are a major focus in regenerative medicine (Shi et al., 2023; Sorg et al., 2017). During the proliferative phase of tissue repair, angiogenesis is initiated in response to local hypoxia, which stabilizes HIF-1α-mediated VEGF signaling pathways. This VEGF upregulation together with other proangiogenic factors such as FGF-2 and PDGF, triggers endothelial activation, basement membrane remodeling, and sprouting vessels from pre-existing vasculature (Cañedo-Dorantes and Cañedo-Ayala, 2019; Carmeliet and Jain, 2011; Veith et al., 2019).

Mesenchymal stem cells (MSCs) are promising candidates for wound healing therapies, not only through their differentiation potential but also due to the secretion of trophic and immunomodulatory factors (Fan et al., 2020; L et al., 2019; Miceli et al., 2021). The angiogenic and regenerative effects of the MSC secretome have been validated in multiple preclinical models, where administration of MSC-derived secretome leads to enhanced wound closure, increased neovascularization, and improved matrix remodeling (Prado-Yupanqui et al., 2025). The MSC secretome contains a broad repertoire of cytokines and growth factors, which can promote angiogenesis and accelerate tissue repair (Liu et al., 2013; Maacha et al., 2020; Prado-Yupanqui et al., 2025). Importantly, secretome composition and potency can be modulated by *in vitro* cell preconditioning strategies that are increasingly explored including pro-inflammatory cytokines or exposure to hypoxia, which augment the release of pro-angiogenic mediators (Joshi et al., 2025; Mrahleh et al., 2021; Oses et al., 2017; Wobma et al., 2018). Cannabinoids, secondary metabolites and bioactive constituents of *Cannabis sativa*, are attracting growing biomedical interest across a range of conditions. In the context of angiogenesis, they have emerged as promising modulators. Among them cannabidiol (CBD), the main nonpsychoactive constituent of *C. sativa*, exhibits antioxidant, anti-inflammatory, and immunomodulatory activity (Atalay et al., 2019; Ujváry and Hanuš, 2016). Nevertheless, although cannabis-based medicines are approved for select neurological indications (Cristino et al., 2020; Devinsky et al., 2024; Manzoni et al., 2025), their potential in tissue repair and regenerative medicine remains insufficiently defined, and the underlying mechanisms of action are poorly characterized. Preclinical evidence suggests that CBD can modulate endothelial function and angiogenesis (Maia et al., 2023). Because MSCs express receptors and channels responsive to cannabinoids (Mesas et al., 2025), we therefore asked whether purified CBD or full-spectrum CBD extracts can stimulate MSCs to enhance their pro-angiogenic secretome. However, because reproducibility and standardization are prerequisites for clinical translation, rigorous phytochemical characterization of full-spectrum cannabis extract is essential to ensure defined composition, batch-to-batch consistency, and predictable bioactivity.

Despite the expanding therapeutic use of cannabinoids, critical evidence gaps remain regarding clinical efficacy, particularly in tissue repair contexts. Preclinical wound healing models suggest pro-regenerative signals may be engaged, yet the mechanistic underpinnings of CBD action remain largely unresolved (Morales et al., 2017). To address this gap, we tested the hypothesis that purified CBD or a standardized full-spectrum CBD-rich Chemotype III (QIII, high CBD:THC ratio) *C. sativa* extract can activate MSC signaling and secretory activity to potentiate angiogenesis, thereby providing mechanistic insight into their potential in tissue repair and regeneration.

Here, we investigated purified CBD alongside a chemically characterized QIII full-spectrum extract in umbilical cord-derived human MSCs (UC-hMSC). We performed phytochemical profiling of the full-spectrum CBD extract to ensure defined composition with the goal of comparing whole full-spectrum extract versus purified CBD effects on UC-hMSC paracrine activity. We focused on transcriptional and secretory responses relevant to angiogenesis, with emphasis on the HIF-1α/VEGF axis. We hypothesize that priming with purified CBD or the QIII full-spectrum extract stabilizes HIF-1α in UC-hMSC, increases the secretion of pro-angiogenic factors, like VEGFA, and yields a secretome with greater angiogenic capacity. Functional consequences of UC-hMSC preconditioning were tested using endothelial proliferation, and tube-formation assays *in vitro*, and vascular branching in the chick chorioallantoic membrane (CAM) model *in vivo*. This integrated approach is designed to clarify the mechanistic basis of cannabinoid-mediated enhancement of MSC pro-angiogenic function and to inform the development of standardized cannabinoid priming for wound healing and regenerative medicine.

## Materials and methods

### Purification and characterization of CBD

Purified cannabidiol (CBD) was obtained from *Cannabis sativa* L. inflorescences (chemotype III) cultivated in a licensed facility in Argentina. Inflorescences were dried, ground, and decarboxylated in a vacuum oven (120°C, 1.5h). Ethanol extraction was performed on decarboxylated biomass using cold ethanol (0°C, 6 h). The crude QIII full-spectrum extract was concentrated under reduced pressure and further purified by two successive flash chromatography steps, yielding a fraction with >99% CBD purity and no traces of other cannabinoids or terpenes, as confirmed by gas chromatography-mass spectrometry (GC-MS).

### Plant material and extraction of phytocannabinoids of Chemotype III full-spectrum CBD extract

A *C. sativa* variety developed by the Interdisciplinary Cannabis Program, CCT CONICET-CENPAT (Puerto Madryn, Argentina), and registered at the National Seed Institute of Argentina (INASE) under the name of “Pachamama” was used for this work. This variety is a chemotype III (QIII) with a CBD:THC > 1. Plants were obtained by clonal propagation, grown on inert substrate and irrigated using an automated system with macro and micro nutrients controlled fertilization. All environmental variables, including temperature, humidity, light intensity, and photoperiod, were monitored during the whole cultivation cycle. No pesticides were used in any stage of the process.

Cannabis dried inflorescences were previously decarboxylated in an oven at 140 °C during 2 h and then extracted at −20 °C for 20 minutes using ethanol 96% at a ratio of 1L per 50 g of dry flowers. Plant material was removed by filtration, and the extract was evaporated in a rotatory evaporator to dry, obtaining a full-spectrum cannabis resin. Major cannabinoids and terpenes were quantified by HPLC-UV and GC-MS, respectively. Cannabis resins were resuspended in DMSO to the working concentrations indicated for each experiment; vehicle controls received an equivalent amount of DMSO.

### Isolation, culture, and identification of human umbilical cord-derived mesenchymal stem cells

Human mesenchymal stem cells were isolated from umbilical cords (UC-hMSC) obtained after full-term deliveries, following informed consent from mothers. All procedures were conducted under protocols approved by the Institutional Research Ethics Committee of Hospital Español de Mendoza, and were carried out for research and potential therapeutic applications.

Umbilical cords were processed preferably within 24 h *postpartum*. Cord segments were placed with the dorsal side facing down on gelatin-coated culture plates, ensuring direct contact between the Wharton’s jelly, rich in stromal hMSCs, and the growth substrate. Explants were incubated in α-MEM supplemented with 20% fetal bovine serum (FBS, SIGMA) at 37°C in a humid atmosphere containing 5% CO₂. The medium was refreshed every 2-3 days. Over a period of 14 to 20 days, hMSCs migrated from the tissue fragments, adhered to the culture surface, and formed colonies.

Once colonies reached approximately 70% confluence, cells were expanded in α-MEM supplemented with 10% FBS, 100 U/mL penicillin, and 100 μg/mL streptomycin, maintaining incubation conditions at 37°C and 5% CO₂. For long-term storage, cells were cryopreserved in the liquid nitrogen atmosphere (−196°C).

Related cell surface markers, including CD29, CD73, CD90, and CD105, were detected by flow cytometry using the BD Stemflow™ Human MSC Analysis Kit (BD Biosciences, Cat. 562245, San Diego, CA, USA); the endothelial marker CD31 was analyzed as a negative marker.

### UC-hMSC preconditioning and secretome collection

For UC-hMSC priming, a purified CBD and full-spectrum CBD QIII extracts were used. All comparisons were performed at CBD-isodose of 0.2 μM (purified CBD vs full-spectrum extract adjusted to the same CBD concentration), thereby controlling for cannabidiol load. UC-hMSC between passages 3 and 6 were seeded to reach ~50% confluence in α-MEM supplemented with 10% FBS, 100 U/mL penicillin, and 100 μg/mL streptomycin. Cells were then washed with phosphate-buffered saline (PBS) and preconditioned for 48 h with CBD or QIII full-spectrum extract, diluted in dimethyl sulfoxide (DMSO, <0.1%) as vehicle, in phenol red-free α-MEM without serum.

After 48 h of preconditioning with CBD or QIII full-spectrum extract, cells were washed with PBS and maintained for an additional 48 h in serum free DMEM to allow secretome collection (Supplementary Fig. S1). In parallel, non-preconditioned or vehicle-treated UC-hMSC, were processed identically to obtain the control secretome. The secretome was centrifuged at 400 × g for 10 min and subsequently at 5000 × g for 20 min at 4 °C to remove cellular debris. The supernatant was then filtered through a 0.45 µm membrane and divided into aliquots, which were stored at −80 °C until use. Additionally, in the indicated experiments concentrated secretome were used. Secretomes were concentrated using Amicon Ultra centrifugal filters (3 kDa MWCO, Millipore). Fifty-μL aliquots were centrifuged at 5,000 × g at 4 °C in sequential cycles, adding additional secretome between spins until the full volume was processed. The concentrated secretomes were transferred to 15 μL Amicon devices for a final concentration step, yielding a final volume of ∼500 µL; and aliquoted (25 µL) and stored at −80 °C. Total protein concentration before and after concentration was measured using a BCA assay according to the manufacturer’s instructions. Unless specified, functional assays were performed with unconcentrated secretome.

### MTS Cell Viability Assay

UC-hMSC were seeded in 96-well plates at a density of 3,000 cells per well in 100 µL of complete medium and allowed to adhere overnight. Cells were then exposed to either purified CBD or the QIII full-spectrum *C. sativa* extract, diluted in DMSO (vehicle) to final concentrations of 0.2 µM. After 48 h of treatment, cell viability was assessed using the MTS Cell Proliferation Assay Kit (Abcam, ab197010) according to the manufacturer’s instructions. Briefly, 20 µL of MTS reagent were added to each well and plates were incubated for 1 h at 37 °C, after which absorbance was measured at 490 nm using a microplate reader. Background absorbance from cell-free wells was subtracted, and data were expressed as percentage of viability relative to DMSO control. Each condition was tested in triplicate.

### RNA extraction and quantitative real-time PCR analysis

Total RNA was extracted from cells using TRIzol™ reagent (Invitrogen, Thermo Fisher Scientific, Cat. No. 15596026) according to the manufacturer’s instructions. RNA quantity and purity were assessed on a NanoDrop™ 2000 spectrophotometer (Thermo Fisher Scientific) by measuring the A260/280 and A260/230 ratios. Complementary DNA (cDNA) was synthesized from 2 µg of total RNA using M-MLV Reverse Transcriptase (Promega, Cat. No. M1705) using oligo(dT)14 primers (IDT) and Recombinant RNasin® Ribonuclease Inhibitor (Promega, Cat. No. N2511), following the manufacturer’s instructions. Quantitative real-time PCR was performed on a StepOnePlus™ Real-Time PCR System (Applied Biosystems, Thermo Fisher Scientific) using 2X Brilliant II SYBR® Green QPCR Master Mix (Agilent Technologies, Cat. No. 600828). Relative mRNA abundance was calculated using the 2^-ΔΔCt method, with GAPDH as the internal reference. Primer sequences are listed in Supplementary Table 1.

### Human umbilical vein endothelial cell (HUVEC) culture

HUVEC were isolated as described by Bannoud et al. (Bannoud et al., 2022) and cultured in Dulbecco’s Modified Eagle Medium (DMEM) supplemented with 10% FBS and 1% penicillin-streptomycin. Cells were maintained at 37°C in a humidified 5% CO₂ atmosphere. Cells were detached using 0.05% trypsin-EDTA.

For functional assays, HUVEC were seeded into the appropriate culture plates and allowed to attach overnight. The next day, culture medium was replaced with preconditioned UC-hMSC-derived secretome for 24 h prior to performing the assays.

HUVEC were pre-treated with the VEGF RTK inhibitor (Calbiochem, cat. 341610; 3 μM in 0.1% DMSO) for 30–60 min, followed by rVEGF165 stimulation (50 ng/mL, 5–10 min, Biotechne R&D System, cat. 293-VE) prior to lysis for p-VEGFR2/p-ERK immunoblotting.

### Tube formation assay

HUVEC were cultured to ∼80% confluence, detached with 0.05% trypsin-EDTA (Gibco, Thermo Fisher Scientific), and seeded into 24-well plates at a density at 2×10⁵ cells per well and incubated at 37°C, 5%CO_2_. After overnight attachment, the medium was replaced with preconditioned UC-hMSC-derived secretome and cells were incubated for 24 h. Subsequently, 10,000 cells from each condition were seeded into 96-well plates pre-coated with 40 μL of growth factor–reduced extracellular matrix (Geltrex, Gibco). Cells were maintained under standard culture conditions (37 °C, 5% CO₂) for 6 h to allow capillary-like tube formation. Tube structures were visualized using a bright-field inverted microscope at 4× magnification. For each well, four random fields were acquired, and all conditions were assayed in triplicate. Vascular endothelial growth factor rVEGF165 (50 ng/ml, Biotechne R&D System, cat. 293-VE) was included as positive control. Quantitative analysis of angiogenic parameters (number of nodes, meshes, junctions, and segments) was performed using the Angiogenesis Analyzer plugin in ImageJ software (NIH, Bethesda, MD, USA).

### Cell proliferation assay (MTS)

HUVEC were seeded into 96-well plates at a density of 5,000 cells per well. After allowing cells to attach, they were treated with preconditioned UC-hMSC-derived secretome for 24 h. Cell proliferation was assessed using the MTS Cell Proliferation Assay Kit (Colorimetric, Abcam, ab197010) according to the manufacturer’s instructions. The MTS reagent was added directly to the wells, and absorbance was measured at 490 nm after 1 h of incubation using a microplate reader. Results are expressed as relative proliferation compared to untreated controls.

### Chick chorioallantoic membrane (CAM) assay

This study was approved by the Scientific Ethics Committee for the Care of Animals and the Environment (CEC-CAA), Pontificia Universidad Católica de Chile (protocol 251014002). Fertilized chicken eggs were obtained from the poultry supplier Chorombo S.A. The assays were performed with adaptations from (Victorelli et al., 2022). On embryonic development day 1 (EDD1), eggshells were disinfected with 70% ethanol and eggs were placed in a humidified incubator at 37.7 °C with gentle automatic rotation (half-turn every 30 min). On EDD3, an approximately 2 cm² window was opened at the thinner end of the egg, and 2–3 mL of albumin was withdrawn using a sterile needle and syringe. The window was then sealed with sterile transparent adhesive tape, and eggs were returned to the incubator.

CAM assays were initiated on ED11, when blood vessels had reached a suitable size for analysis (>80 µm in diameter). For treatment, 100 µL of preconditioned UC-hMSC-derived secretome was applied either unconcentrated or concentrated to 10 μg/mL directly onto the CAM. Eggs were randomly assigned to treatment groups. The CAM was imaged at baseline (0 h) and again at 24, 48, and 72 h after treatment using a digital USB microscope. For the control group, the images were performed at the same time in the CAM without any application.

Vascular branching points were quantified from digital images using ImageJ software, considering nodes comprising at least three branches within a predefined region of interest. All procedures were performed under aseptic conditions in a laminar-flow cabinet.

### SDS-PAGE and Western blot analysis

UC-hMSC were lysed in RIPA buffer supplemented with protease (P2714, Sigma-Aldrich) and phosphatase inhibitors (P2714, Sigma-Aldrich). Lysates were incubated on ice for 15 min with vortexing every 5 min, and then centrifuged at 12,000 × g for 15 min at 4 °C. The resulting supernatants were collected and stored at −20° C until analysis. Protein concentration was determined using the BCA Protein Assay Kit (Thermo Fisher Scientific), and 25–30 µg of total protein was loaded per lane. Proteins were resolved by SDS-PAGE (10%) and transferred to 0.45 µm nitrocellulose membranes (Bio-Rad).

Membranes were blocked in 5% BSA in TBST (Tris-buffered saline, 0.1% Tween-20) for 1 h at room temperature and incubated overnight at 4°C with the primary antibodies (see Supplementary Table 2). After three washes in TBST, membranes were incubated with HRP-conjugated secondary antibodies (see Supplementary Table 2) for 1 h at room temperature, washed again, and developed by enhanced chemiluminescence (ECL, Kalium Technologies, Buenos Aires, Argentina). Images were acquired on a LAS-4000 luminescent imager. When densitometry was performed, band intensities were quantified using Fiji (version 1.54), normalized to α/β-tubulin (loading control) or total ERK (for phospho/total ratios), and expressed relative to the indicated control condition.

### Statistical analyses

Statistical analyses were performed using R with RStudio (version 2025.05.0+496) and the packages *tidyverse*, *ggpubr*, and *dplyr*. Data were first tested for normality using the Shapiro-Wilk test and for homogeneity of variances using Bartlett’s test. When assumptions were met, group means were compared using one-way analysis of variance (ANOVA), followed by Tukey’s or Dunnett’s post hoc tests when appropriate. For the CAM assay the two-way ANOVA was performed, followed by Tukey’s post-hoc multiple comparison test. A p<0.05 was considered statistically significant. Graphical representation of the data was generated in Prism 9 (version 9.5.1).

## Results

### Phytochemical characterization of the full-spectrum CBD Chemotype III *Cannabis sativa* extract

To ensure reproducibility and define the composition of the plant material used in this study, we first characterized the phytochemical profile of the full-spectrum CBD *C. sativa* extract (QIII). HPLC-UV chromatographic analysis revealed distinct peaks corresponding to major cannabinoids, including cannabidiol (CBD), cannabidiolic acid (CBDA), Δ^9^-tetrahydrocannabinol (THC), tetrahydrocannabinolic acid (THCA), and cannabinol (CBN) (Fig. 1A). Quantitative analysis confirmed that CBD was the predominant constituent 52.7 mg/mL, followed by CBDA (2.0 mg/mL), THCA (2.2 mg/mL), THC (1.8 mg/mL), and CBN (0.9 mg/mL) (Fig. 1B). The resulting CBD:THC ratio (11.2:1) was consistent with the chemotype III classification. The data established a well-defined cannabinoid composition for the QIII extracts, providing a reproducible chemical profile for subsequent experiments.

**Fig. 1.**
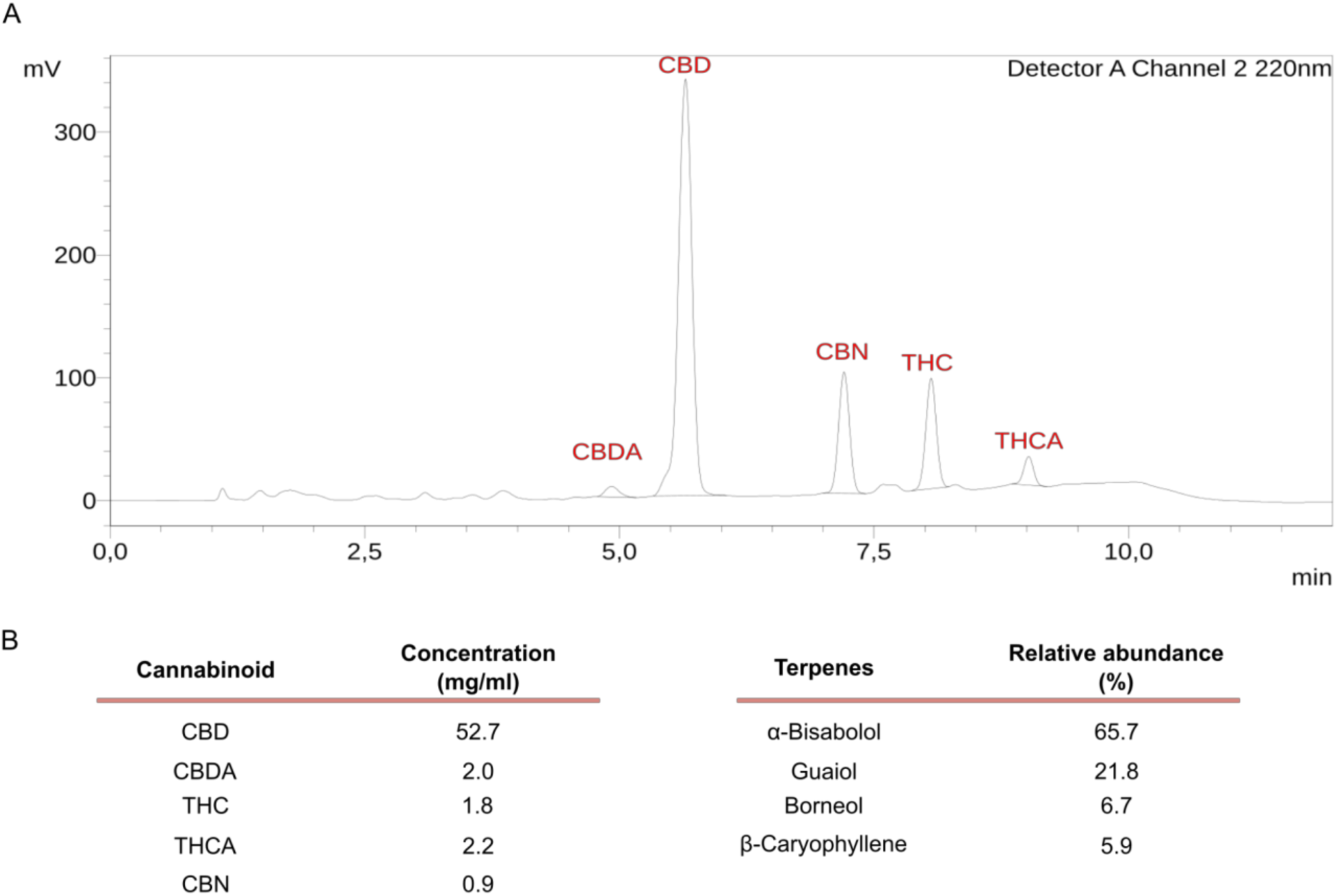
Phytochemical characterization of the full-spectrum CBD Chemotype III *Cannabis sativa* extract. (**A**) HPLC-UV chromatogram of the cannabinoid fraction with peak assignments for CBD, CBDA, THC, THCA, and CBN. (**B**) Quantification of major cannabinoids in the ethanolic extract (mg/mL) by HPLC-UV. (**C**) Relative abundance (%) of principal terpenes by GC-MS.

We then analyzed the volatile fraction by GC-MS. The chromatographic profile revealed the presence of mono- and sesquiterpenes. Identification was carried out by comparison with a commercial terpene standard mix (Terpenes MegaMix#1, Restek), and the relative abundance of each compound. The extract was dominated by sesquiterpenes, with α-bisabolol (65.7% of total identified terpenes) and guaiol (21.8%) representing the most abundant constituents. Other relevant components included borneol (6.7%) and β-caryophyllene (5.9%) (Fig. 1C).

### Purified CBD and full-spectrum CBD extract modulate the transcriptome of UC-hMSC toward a pro-angiogenic and stress-resilient phenotype

We first verified UC-hMSC identity by flow cytometry (Fig. 2A). Preconditioning at the indicated doses did not compromise viability(Supplementary Fig. S2); values were normalized to the vehicle and interpreted according to ISO 10993-5 (≥ 70% non-cytotoxic).

**Fig. 2.**
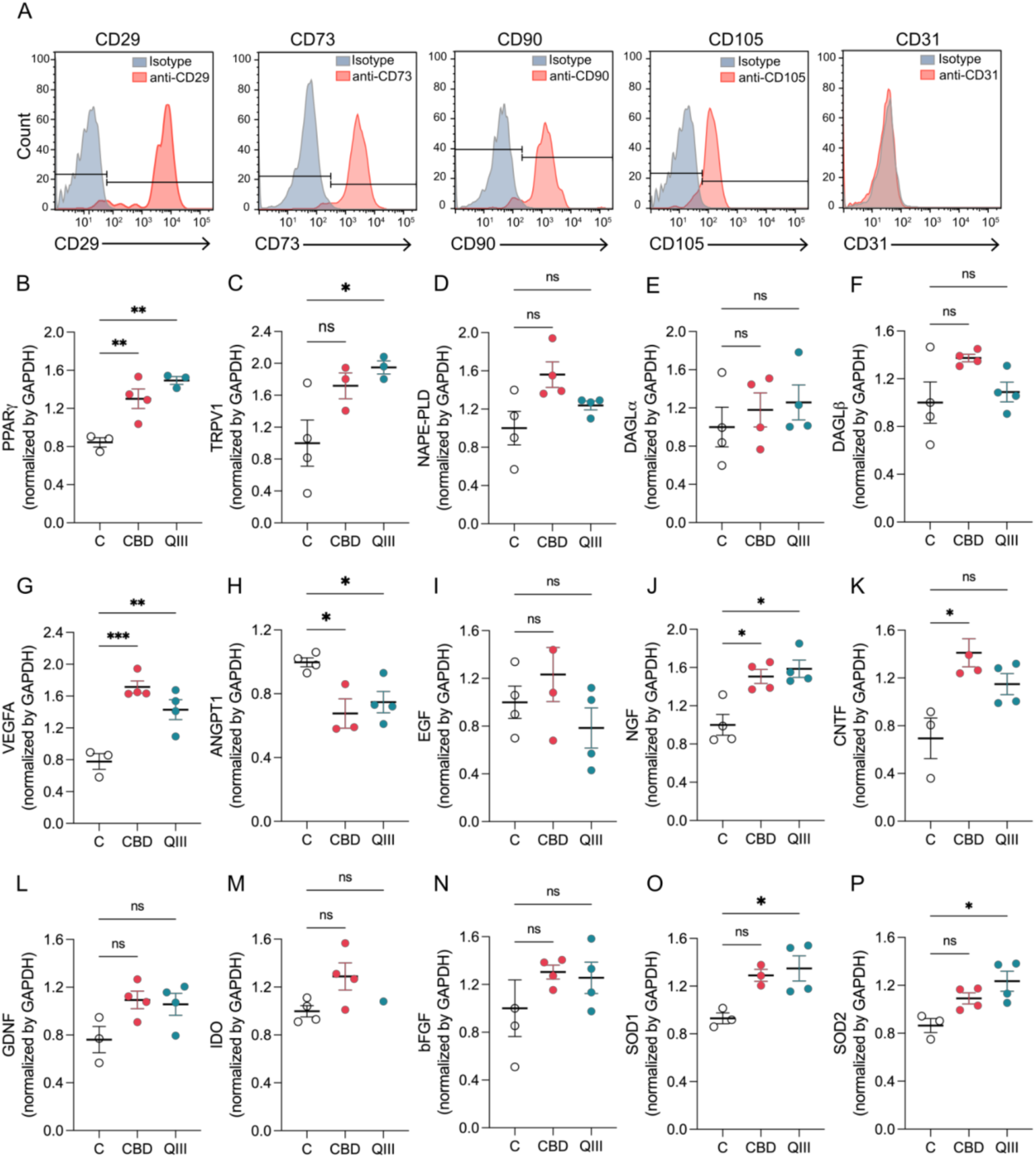
Purified CBD and full-spectrum CBD extract (QIII) priming of UC-hMSC activates transcriptional profiles relevant to angiogenesis. (**A**) Characterization of UC-hMSC. Flow cytometry staining demonstrated the negative expression of endothelial marker (CD31), and were positive for MSC markers (CD29, CD73, CD90 and CD105). Quantitative RT-qPCR analysis in UC-hMSC exposed to vehicle (C), purified CBD (CBD), or a QIII full-spectrum extract (QIII). Transcriptional levels of (**B**) PPARγ, (**C**) TRPV-1, (**D**) NAPE-PLD, (**E**) DAGLα, (**F**) DAGLβ (**G**) VEGFA, (**H**) ANGPT1, (**I**) EGF, (**J**) NGF, (**K**) CNTF, (**L**) GDNF, (**M**) IDO, (**N**) bFGF, (**O**) SOD1, and (**P**) SOD2 were quantified. Data are shown as mean ± SEM from one representative experiment (n=3), with each dot representing a technical replicate. Data are represented as mean ± SEM. Statistical significance was determined using one-way ANOVA, *p < 0.05, **p < 0.01, ***p < 0.001, ns = not significant.

We then asked whether cannabinoid priming engages gene programs in MSC that are compatible with a pro-angiogenic secretome. We analyzed the expression of two non-canonical CBD targets and endocannabinoid system (ECS) components. RT-qPCR revealed that mRNA PPARγ was significantly upregulated after purified CBD and QIII full-spectrum extracts preconditioning, while TRPV1 showed a significant increase with QIII full-spectrum priming (Fig. 2B, C). Given that cannabinoids can also act indirectly on the ECS, we examined biosynthetic enzymes controlling anandamide (AEA) and 2-arachidonoylglycerol (2-AG) production. Neither NAPE-PLD (which catalyzes the synthesis of AEA), and DALGα/β (responsible for generating 2-AG) transcripts were significantly altered, although we observed a modest increase in purified CBD-preconditioned UC-hMSC (Fig. 2D, E, F).

We next evaluated angiogenesis-related genes. VEGFA was robustly induced by both purified CBD and QIII full-spectrum extracts, whereas ANGPT1 showed significant reduction in both stimuli, consistent with a shift toward immature sprouting neovasculature (Fig 2G, H). EGF expression also tended to increase in purified CBD-preconditioned UC-hMSC (Fig 2I). We then examined transcripts linked to immunomodulation and trophic support that can influence tissue repair and angiogenesis. In addition, these mediators were differentially affected, NGF increased significantly with both treatments, CNTF augmented with purified CBD, and GDNF remained unchanged (Fig 2J, K, L). IDO did not differ significantly, although showed an increase with QIII full-spectrum extract, whereas bFGF was modestly enhanced in both conditions (Fig. 2M, N). Finally, to probe whether CBD-driven activates redox control, we examined antioxidant defenses and found that SOD1 and SOD2 were significantly upregulated by QIII full-spectrum extract, with a non-statistically significant increase under purified CBD (Fig 2O, P). This coordinated induction suggests reinforcement of cytosolic and mitochondrial antioxidant capacity, with a stronger effect following QIII full-spectrum extract preconditioning.

Taken together, these data suggest that purified CBD and QIII full-spectrum extracts activates UC-hMSC by inducing PPARγ and selectively enhancing VEGFA, NGF, and CNTF, while modestly affecting ECS enzymes and classic immunomodulators. This transcriptional profile is complemented by increased SOD1/2 expression, consistent with strengthened antioxidant defenses. Collectively, cannabinoid preconditioning endows MSC with a pro-angiogenic, trophic, and stress resilient phenotype, prompting us to evaluate the functional impact of their secretomes on endothelial cells.

### Cannabinoid preconditioning activates ERK and upregulates VEGFA in UC-hMSC

To assess early signaling events associated with transcriptional activation, we examined ERK1/2 phosphorylation by immunoblotting in UC-hMSC after preconditioning for 30 minutes. Both purified CBD and QIII full-spectrum extracts increased phospho-ERK (p-ERK) relative to total ERK. Densitometric analysis confirmed a significant rise in p-ERK/ERK ratios (Fig. 3A). These data together with VEGFA mRNA induction (Fig. 1G) indicate engagement of the ERK pathway by cannabinoid preconditioning. We then asked whether the transcriptional induction of VEGFA is reflected at the protein level in UC-hMSC lysates. At 24 h, immunoblotting showed a significant increment in intracellular VEGFA with CBD preconditioning (Fig. 3B). By 48 h, both purified CBD and the QIII full-spectrum extracts showed lower intracellular VEGFA than control (Fig. 3C). In the context of early ERK activation and VEGFA mRNA and protein increase, the reduction of VEGFA at 48 h in cell lysates may reflect increased secretion into the secretome.

**Fig. 3.**
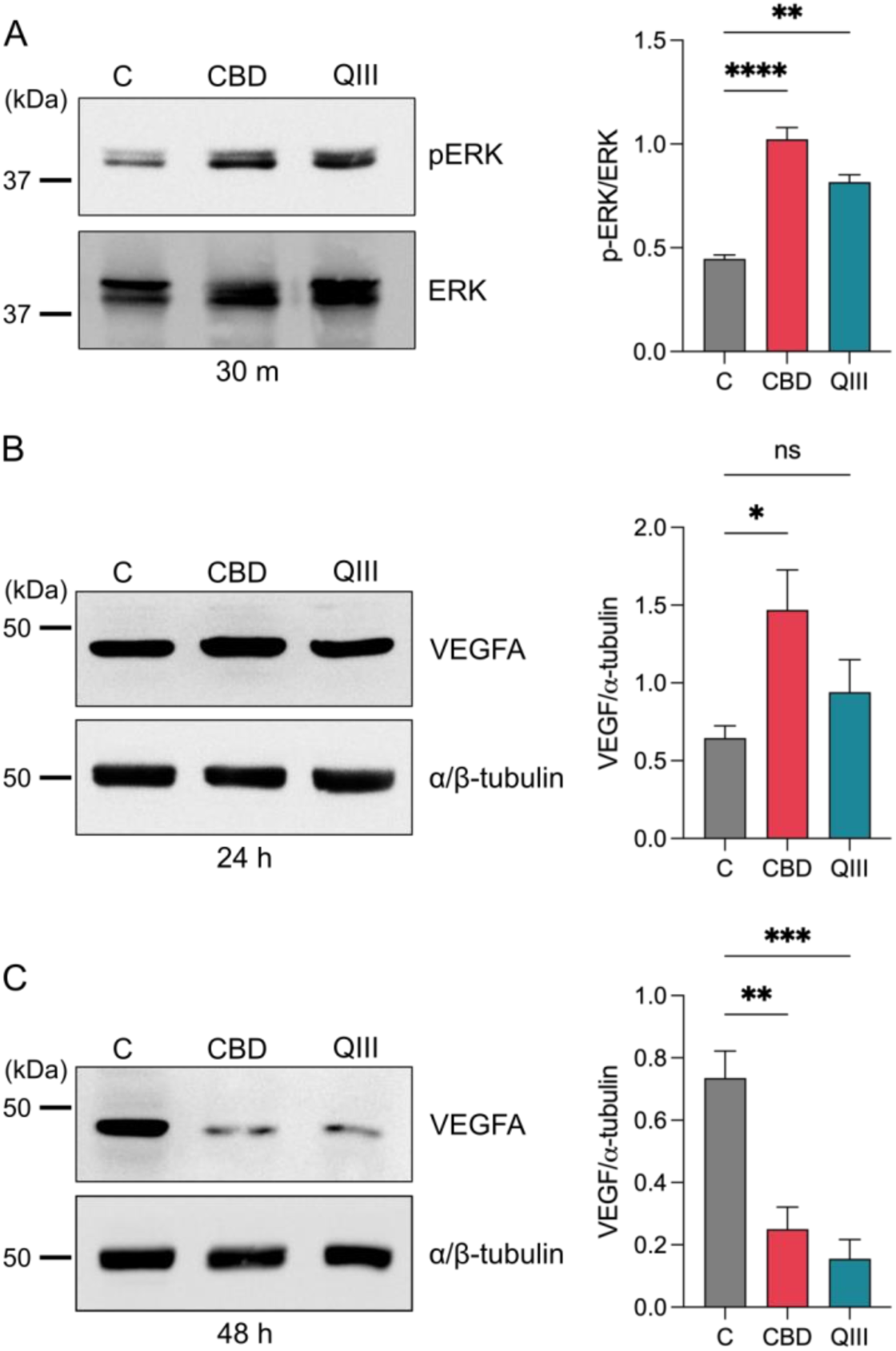
Early ERK activation and VEGFA redistribution in UC-hMSC after purified CBD or QIII full-spectrum preconditioning. (**A**) Representative immunoblot of p-ERK1/2 and total ERK in UC-hMSC after 30 min, unprimed UC-hMSC (C), purified CBD (CBD), or QIII full-spectrum extract (QIII) preconditioning. Densitometry of p-ERK/ERK (n=3). (**B**) Representative immunoblot of VEGFA in UC-hMSC lysates following 24 h of treatment, control (C), CBD (CBD), or QIII extract (QIII). α/β-tubulin is shown as loading control (n=3). Densitometry of VEGFA normalized to α/β-tubulin. (**C**) Representative immunoblot of VEGFA in UC-hMSC lysates following 48 h of treatment control (C), purified CBD (CBD), or QIII full-spectrum extract (QIII). α/β-tubulin is shown as loading control. Densitometry of VEGFA normalized to α/β-tubulin (n=4). Data are represented as mean ± SEM. Statistical significance was determined using one-way ANOVA, *p < 0.05, **p < 0.01, ***p < 0.001, ****p < 0.0001, ns = not significant.

### Purified CBD and QIII full-spectrum extracts preconditioning of UC-hMSC enhances the pro-angiogenic activity of their secretome *in vitro*

After showing that purified CBD and QIII full-spectrum extracts induce activation of UC-hMSC toward a pro-angiogenic and antioxidant phenotype, we next assess whether their secretomes exert functional effects on endothelial cells. Treatment of HUVEC with CBD (sCBD) and QIII extract (sQIII) primed secretomes significantly enhanced proliferation at 24 h compared with secretome from unprimed UC-hMSC (sC), with the strongest effect observed in the QIII extract group (Fig. 4A). We next investigated their effects on capillary-like tube formation. HUVEC exposed to secretomes derived from UC-hMSC preconditioned with purified CBD or the QIII full-spectrum extract formed more complex networks than control (Fig. 4B). Quantification showed that the number of nodes, junctions and segments, were significantly higher in both conditions, whereas the number of meshes was significantly increased with QIII extract (Fig. 4C-F). Together, these data indicate that purified CBD and QIII full-spectrum extracts preconditioning endows UC-hMSC secretomes with enhanced angiogenic activity *in vitro*.

**Fig. 4.**
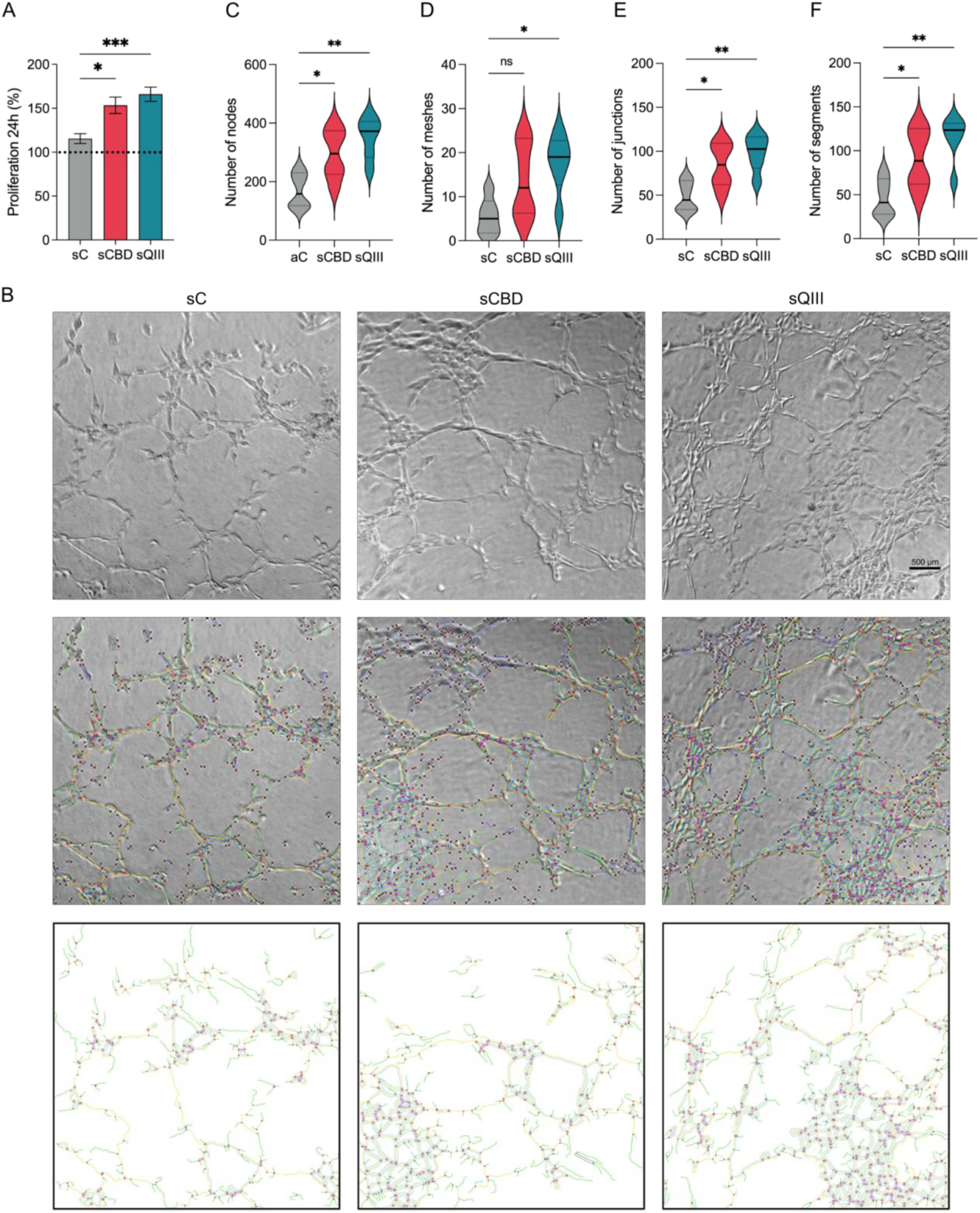
Purified CBD and full-spectrum extract (QIII) primed UC-hMSC secretomes promoted HUVEC proliferation and tube formation *in vitro*. (**A**) HUVEC proliferation after 24 h of treatment with unprimed UC-hMSC (sC), CBD-preconditioned UC-hMSC (sCBD), or QIII extract-preconditioned UC-hMSC (sQIII). (**B**) Tube formation assay, representative images after being treated with the indicated secretomes as in (A). Violin plots show quantification of (**C**) number of nodes, (**D**) number of meshes, (**E**) number of junctions, and (**F**) number of segments. Data are shown as mean ± SEM, n=3 biological replicates, each technical replicate has been plotted. Scale bar: 500 μm. Statistical significance was determined using one-way ANOVA followed by post hoc test, *p < 0.05, **p < 0.01, ***p < 0.001, ns = not significant.

### Secretomes from purified CBD and QIII full-spectrum extracts preconditioned UC-hMSC increase vascular branching *in vivo*

We next validated angiogenic activity in a physiological setting, we used the chick chorioallantoic membrane (CAM) assay, a well-established model for assessing neovascularization.

Representative images obtained 24 h after secretome application, revealed increased vascular branching in embryos treated with purified CBD or QIII full-spectrum extract primed secretomes compared with control secretome for untreated condition (Fig. 5A). The images clearly show an increase in vessel density 24 h after the application of CBD and QIII full-spectrum extracts. Quantitative analysis confirmed that the number of branching points was significantly higher in both groups (Fig. 5B). An extended time course showed that this response was less marked at 48 h and declined to baseline at 72 h. Under concentrated conditions, the increase was only observed at the longer time point (72 h), particularly for CBD (Supplementary Fig. S3). These findings demonstrate that CBD and QIII extract preconditioning enhances the capacity of UC-hMSC secretomes to stimulate neovascular sprouting *in vivo*, in agreement with the *in vitro* data observed in HUVEC cells, supporting the view that cannabinoid priming strengthens the pro-angiogenic capacity of the UC-hMSC secretome.

**Fig. 5.**
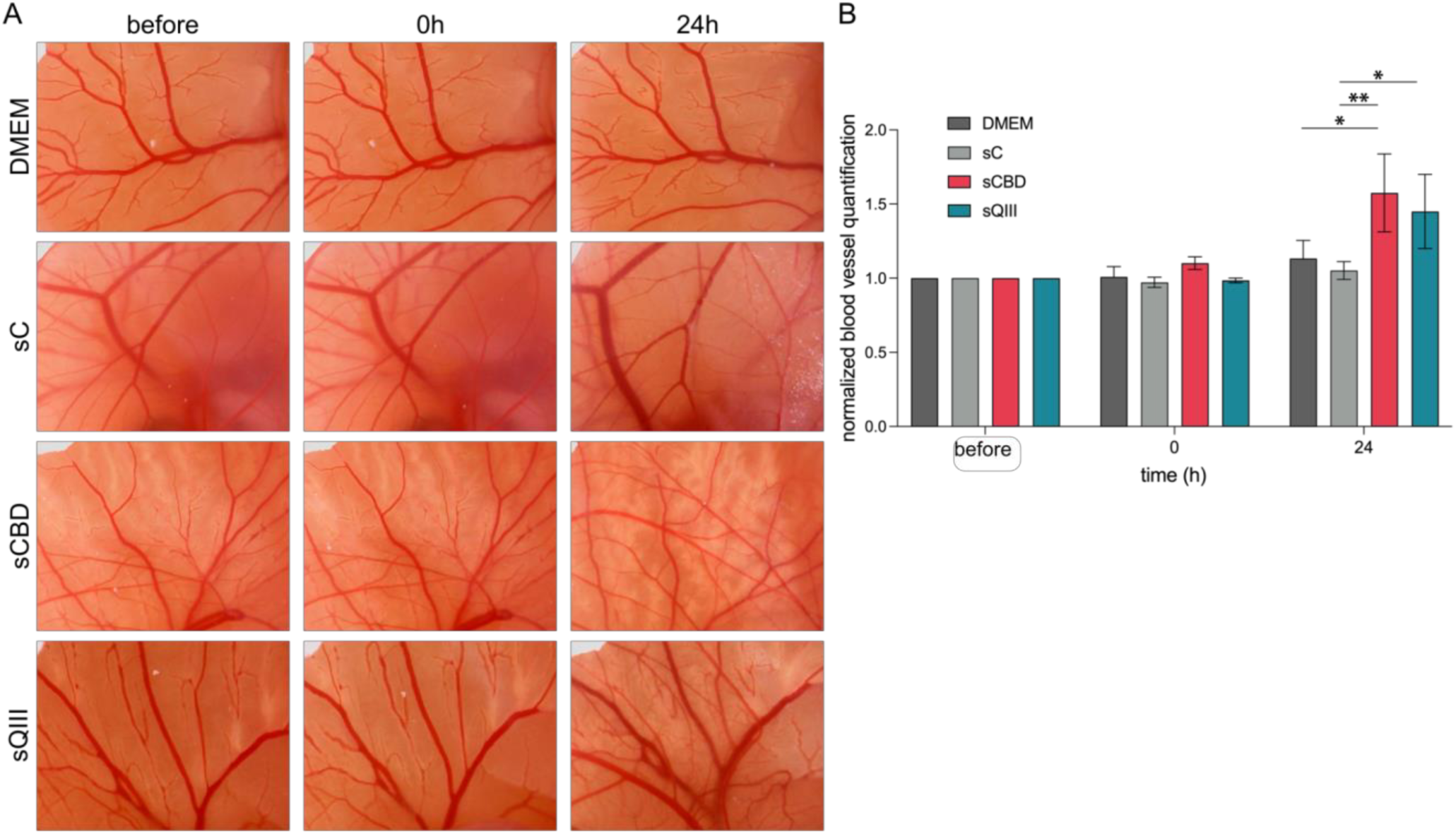
Purified CBD and QIII full-spectrum extracts primed UC-hMSC secretomes enhance angiogenesis in the chick chorioallantoic membrane (CAM) assay. (**A**) Representative images of the CAM assay at 0 h and 24 h after applying DMEM, and conditions for unprimed UC-hMSC secretome (sC), purified CBD-primed UC-hMSC secretome (sCBD), and QIII full-spectrum extracts-primed UC-hMSC secretome (sQIII). (**B**) Quantification of vascular branching points. Data are shown as mean ± SEM from n= 8 eggs per group (biological replicates). Statistical analysis was performed by one-way ANOVA followed by Tukey’s post hoc test, *p < 0.05, **p < 0.01.

### Secretomes from purified CBD and QIII full-spectrum extracts primed UC-hMSC activate HIF-1α/VEGF–VEGFR-2 signaling

After confirming *in vitro* and *in vivo* that secretomes from CBD and QIII extract primed UC-hMSC increased vascular branching in the CAM assay, we next investigated the underlying mechanism, linking priming of UC-hMSC to endothelial responses.

In UC-hMSC, both purified CBD and the QIII full-spectrum extract stabilized HIF-1α (Fig. 6A). HIF-1α provides a plausible upstream hub integrating redox and metabolic cues to drive a pro-angiogenic transcriptional program in UC-hMSC, with VEGFA as a canonical effector. In HUVEC cells exposed to secretome from unprimed UC-hMSC (sC), purified CBD preconditioned UC-hMSC (sCBD), or QIII full-spectrum extract preconditioned UC-hMSC (sQIII), sCBD significantly increased the phosphorylated ERK signal, whereas sQIII did not differ significantly from control (sC) (Fig. 6B). These data indicate that purified CBD preconditioning enhances the capacity of the secretome to activate ERK signaling in endothelial cells. We next analyzed if the activation of ERK is driven by VEGF-VEGFR2 signaling, HUVEC were exposed to the secretomes obtained as mentioned before in the presence of a VEGFR-2 tyrosine-kinase inhibitor. Pharmacological blockade reduced ERK1/2 phosphorylation across all conditions, lowering the values observed in the absence of the inhibitor (Fig. 6B), indicating that ERK activation by the secretome is VEGFR-2 dependent, in agreement with VEGFA acting as a principal upstream driver (Fig. 2G).

**Fig. 6.**
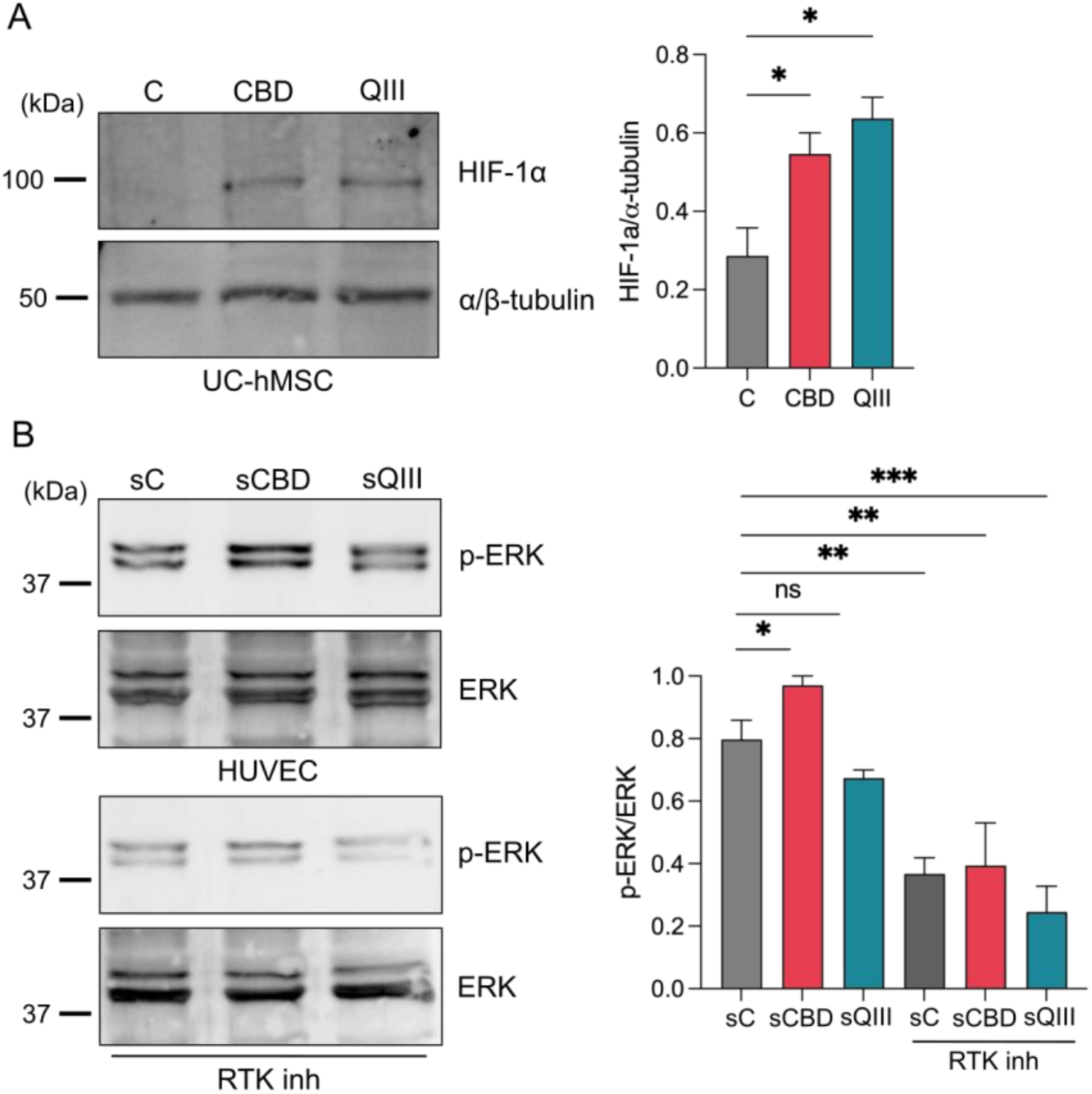
Cannabinoid priming stabilizes HIF-1α in UC-hMSC and primed-secretome drives VEGFR-2-dependent ERK activation in HUVEC. (**A**) Representative immunoblot of HIF-1α in UC-hMSC after 48 h, control (C), purified CBD (CBD), or QIII full-spectrum extract (QIII) preconditioning. α/β-tubulin is shown as loading control. Densitometry of HIF-1α normalized to α/β-tubulin (n=3). (**B**) Representative immunoblot of phospho-ERK1/2 and total ERK in HUVEC cells following 24 h of treatment with unprimed UC-hMSC secretome (sC), purified CBD-primed UC-hMSC secretome (sCBD), and QIII full-spectrum extract-primed UC-hMSC secretome (sQIII) in the absence or presence of RTK inhibitor. Densitometry of p-ERK/ERK (n=3). Data are represented as mean ± SEM. Statistical significance was determined using one-way ANOVA followed by Dunnett’s post hoc test, *p < 0.05, **p < 0.01, ***p < 0.001, ns = not significant.

Taken together, these findings support a model in which CBD and the QIII full-spectrum extract preconditioning of UC-hMSC modulates their transcriptional levels leading to upregulation of pro-angiogenic and antioxidant genes, stabilizing HIF-1α, and inducing VEGFA expression. These molecular changes translate into secretome with enhanced angiogenic activity, as evidenced by increased HUVEC proliferation, and tube formation, as well as augmented neovascular branching in the CAM assay. Notably, endothelial ERK phosphorylation was consistently elevated with purified CBD preconditioned secretome, whereas QIII full-spectrum behaved similarly to control, indicating that VEGFR-2-dependent ERK engagement is prominent for purified CBD and that QIIII-driven effects rely on additional or ERK-independent signaling.

## Discussion

Impaired angiogenesis is a central barrier to chronic wound repair, and multiple strategies to have been studied to potentiate the paracrine angiogenic output of MSC, including hypoxia, cytokine exposure, and pharmacological priming (González-Rodríguez et al., 2025; Noronha et al., 2019; Yang et al., 2022).

While cannabidiol (CBD) and *Cannabis sativa* preparations have been examined across diverse biological contexts, whether these compounds reprogram the paracrine output of MSCs remains underexplored (Rossi et al., 2013; Xie et al., 2016).

Here, we show that preconditioning of UC-hMSC with either purified CBD or a chemically standardized Chemotype III full-spectrum *C. sativa* extract enhances a pro-angiogenic secretome capable of promoting endothelial responses *in vitro* and *in vivo*. We showed that both preconditioning activated HIF-1α/VEGFA signaling in UC-hMSC, providing a mechanistic basis for the VEGFR-2/ERK-dependent endothelial activation observed.

At the transcriptional level, we found that UC-hMSC preconditioned with purified CBD or the QIII full-spectrum extract showed increased expression of VEGFA, NGF, and CNTF, and SOD1/2, consistent with reinforced antioxidant defenses. The corresponding secretomes improved HUVEC proliferation and tube formation, and in the CAM assay increased vascular branching.

Our findings point to a redox, metabolic axis in UC-hMSC in which purified CBD or the QIII full-spectrum extract induces PPARγ, promotes oxidative stress resilience, stabilizes HIF-1α, and increases VEGFA expression. The decline in intracellular VEGFA at 48 h further suggests augmented secretion. Additionally, we observed early ERK1/2 activation in UC-hMSC, followed by VEGFA upregulation, a sequence compatible with ERK-linked transcriptional control. However, whether ERK1/2 is necessary or sufficient for VEGFA upregulation will require inhibition studies during UC-hMSC preconditioning.

Our findings suggest the involvement of at least two upstream mechanisms that likely contribute to VEGFA regulation in UC-hMSC. First, although CBD has low affinity for CB1 and CB2 receptors, it exhibits a broad pharmacological profile that includes interactions with nuclear receptors and ion channels (Castillo-Arellano et al., 2023; Ibeas Bih et al., 2015; Pertwee, 2008). Notably, CBD acts as a well-characterized PPARγ agonist, a nuclear receptor implicated in the modulation of inflammatory and angiogenic responses (Khosropoor et al., 2023). In this sense, we observed an upregulation of PPARγ mRNA levels following cannabinoid preconditioning. This transcriptional response provides a plausible molecular entry point into reparative programs, including redox control and NF-κB transrepression (Dovinova et al., 2020; Varga and Nagy, 2008). These changes could indirectly facilitate the HIF-1α stabilization under stress by shaping the intracellular redox state. Secondly, cannabinoid preconditioning led to increased VEGFA expression in UC-hMSC accompanied by higher HIF-1α levels. This aligns with previous reports indicating that CBD mitigates oxidative stress and regulates the expression of HIF-regulated genes *in vivo* (Kletkiewicz et al., 2024; Yu et al., 2023). Moreover, ERK activity, elevated in our system, can also promote both translation and activity of HIF-1α, suggesting an additional and complementary regulatory pathway. Given that HIF-1α activity is strongly dependent on cellular context and temporal dynamics, its role in our model is likely cooperative, acting in concert with PPARγ-mediated transcriptional reprogramming rather than functioning as an isolated driver.

The predominance of paracrine mechanisms in MSCs mediated repair is well established, and our findings extend this paradigm by adding cannabinoid preconditioning. Secretome derived from purified CBD primed UC-hMSC activated VEGFR-2 and ERK signaling in HUVEC cells, mirroring canonical VEGFA-dependent signaling (Liu et al., 2023). In this sense, VEGFR-2 inhibition, reduced p-ERK across conditions, which is consistent with the principal signaling pathway mediating VEGF-driven angiogenic, and with the functional behavior seen in the endothelial assays (da Rocha-Azevedo et al., 2020).

While CBD-primed secretome increased ERK phosphorylation, this did not translate into the strongest angiogenic effect. This is consistent with reports showing that ERK activation alone does not predict functional angiogenesis, which relies on multiple coordinated pathways (Bullard et al., 2003). In contrast, the QIII full-spectrum (CBD-rich) extract, being a more complex phytochemical mixture, may instead enhance additional pro-angiogenic mediators (e.g., PI3K/AKT) (Chen et al., 2005; Chung et al., 2010; Song and Finley, 2020), explaining its stronger functional effects despite modest ERK activation in HUVEC cells, suggesting alternative angiogenic drivers.

Across several readouts, the QIII full-spectrum extract produced stronger pro-angiogenic effects that purified CBD. Given that the extract is enriched in oxygenated sesquiterpenes (e.g. α-bisabolol and guaiol), with antioxidant and immunomodulatory properties (da Silveira e Sá et al., 2015; Karami et al., 2024), synergy is congruent and aligns with the proposed entourage effect (Russo, 2011). Pharmacological strategies to stabilize HIF-1α have been only slightly explored (Saber et al., 2024; Yuan et al., 2024); in this context, our data point to the QIII full-spectrum extract as a potential stabilizing agent in UC-hMSC. In this sense, the Cannabis medicinal variety developed by CONICET is a good candidate to perform standardized full-spectrum extracts. Nonetheless, disentangling CBD from minor phytocannabinoids and terpenes remains essential for standardization, mechanistic precision, and dose finding in translational settings.

Under oxidative stress, redox sensitive mechanisms promote PPARγ nuclear translocation and amplify its transcriptional activity (Teratani et al., 2020). In this context, the rise in PPARγ that we observed in UC-hMSC following preconditioning offers a coherent rationale for the activation of the HIF-1α/VEGFA axis. In light of evidence that human MSC are endocannabinoid-responsive and that cannabinoid inputs converge on ERK/MAPK pathways (Song and Finley, 2020), our findings support a two-step mechanism in which purified CBD or the QIII full-spectrum extract priming UC-hMSC induce transcriptional and redos remodeling, recalibrates the secretome, activating endothelial VEGFR-2→ERK signaling to drive angiogenic behavior (Fig. 7).

**Fig. 7.**
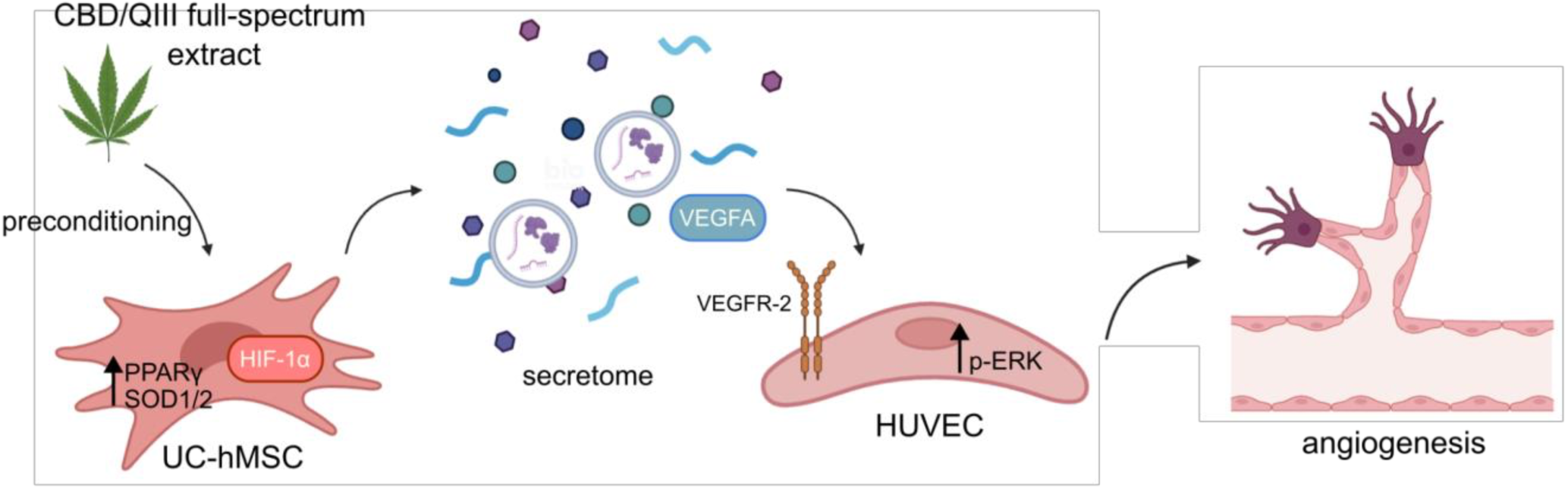
Proposed mechanistic model of cannabinoid preconditioning of UC-hMSC and endothelial readout. Purified CBD and QIII full-spectrum *Cannabis sativa* extract preconditioning of UC-hMSC, induces PPARγ and reinforcing antioxidant defenses (SOD1/SOD2), with stabilization of HIF-1α and increased VEGFA transcription. The resulting pro-angiogenic secretome activates endothelial VEGFR-2, leading to ERK phosphorylation and angiogenic responses—proliferation, capillary-like tube formation, and increased branching in the CAM assay (Created in Biorender.com).

Overall, our results support a working model in which cannabinoid priming promotes angiogenesis via activation of the HIF-1α/VEGF/VEGFR2 axis, with ERK acting as a key mediator in endothelial cells (Fig. 7). In chronic wound contexts characterized by oxidative stress and impaired vascularization, cannabinoid primed MSC secretomes may offer a cell-free therapeutic strategy to restore angiogenic competence and enhance tissue repair.

## Conclusion

Our results show that purified CBD and QIII full-spectrum (CBD-rich) *C. sativa* extracts shift UC-hMSC toward a pro-angiogenic and stress-resilient phenotype. Both treatments increased HIF-1α and VEGFA levels in UC-hMSC and generated a secretome capable of activating endothelial VEGFR-2 and downstream ERK signaling. Together, these findings highlight cannabinoids as modulators of hMSC paracrine activity and emphasize the relevance of comparing purified compounds with full-spectrum preparations when designing translational approaches aimed at restoring angiogenesis in wound healing and regenerative medicine. More broadly, our work supports the growing evidence that the UC-hMSC secretome may serve as a cell-free strategy to promote angiogenic responses. The concept of using phytochemical preconditioning to modulate MSC paracrine outputs represents a promising direction for future studies.

## Limitations and future directions

Regarding the endocannabinoid system, we observed directional non-significant increases in NAPE-PLD and DAGLα/β transcripts, suggesting a potential rise in autocrine endocannabinoid tone; direct lipidomic quantification of AEA and 2-AG, together with pharmacological or genetic manipulation of FAAH and MAGL, will be needed to test causality. PPARγ remains a candidate driver; selective antagonism or genetic interference should be used to determine necessity/sufficiency and its interaction with HIF-1α and ERK/PI3K pathways.

In endothelial cells, VEGFR-2 signaling was clearly engaged; however the degree of VEGF dependence underlying the pro-angiogenic phenotype should be further defined. This should include neutralizing anti-VEGF and/or VEGFR-2 inhibitors across functional assays, as well as quantification of additional secretome components, such as FGF-2, ANGPTL4, and IL-8/CXCL8, that may cooperate with VEGF. Another limitation is the absence of comprehensive secretome profiling. Unbiased proteomics coupled to extracellular-vesicle analysis, and multiplex cytokine quantification will be required to identify mediators modulated by cannabinoid preconditioning and to prioritize targets for functional validation. Finally, vascular maturation was not directly assessed; given the ANGPT1 decrease (trend), systematic evaluation of stabilization markers will help determine whether this shift affects vessel maturation.

Addressing these points will sharpen causal links between specific receptors, endogenous ligands, and defined phytochemical constituents, and will be essential for standardization and future translational applications.

## Author agreement

The authors agree to be responsible for all aspects of the work

## Declaration of competing interests

All authors declare that there are no interests that need to be disclosed in this work.

## Acknowledgments

We thank Dr. Manuel Varas-Godoy (Centro de Biología Celular y Biomedicina, Universidad San Sebastián, Santiago, Chile) for generously providing reagents used in this study. We also thank Gabriel Vaccaro, a solidarity grower registered under Registro del Programa de Cannabis (Ministerio de Salud, Argentina), for providing us with two cannabinoid inflorescences variants for the CBD purification. Mariana Lozada, Tomás Bosco and Rodrigo Barrera helped with cultivation, preparation of full spectrum extracts and quality control. We also thank Fresia Solis for her support in the CAM experiments and the Avícola Chorombo S.A. for their donation of fertile eggs.

This work was supported by Programa de Investigación y Desarrollo en Cannabis, Ministerio Nacional de Ciencia y Tecnología, Argentina (Proyecto 4A) and Agencia Nacional de Promoción Científica y Tecnológica, Argentina (PICT-00668) to M.C.R and (PICT 2019-0532 & PICT 2022-0301) to D.O.C; by Consejo Nacional de Investigaciones Científicas y Técnicas, Argentina (CONICET-PIP-0514) to D.O.C and M.C.R.; by National Institutes of Health (NIH-NCI)-U54 Program CA221208 to D.O.C; by Agencia Nacional de Investigación y Desarrollo, Chile (FONDEF-ID24I10119 and Fondecyt 1250728) to M.E., by Agencia Nacional de Investigación y Desarrollo, Chile (Fondecyt Regular 1241088) to H.H.B, and Programa de Investigación y Desarrollo en Cannabis, Ministerio Nacional de Ciencia y Tecnología, Argentina (Proyecto 6A) to G.B.

## Declaration of Generative AI and AI-assisted technologies in the writing process

During the preparation of this work the author(s) used ChatGPT by OpenAI in order to help with language-related aspects. After using this tool/service, the author(s) reviewed and edited the content as needed and take(s) full responsibility for the content of the publication.

## Notes

### Competing Interest Statement

The authors have declared no competing interest.

