## Supplementary figures and images for "Purified CBD and CBD-rich full-spectrum *Cannabis sativa* extract potentiate the angiogenic paracrine function of umbilical cord derived mesenchymal stem cells"

### Supplemental Figure 1

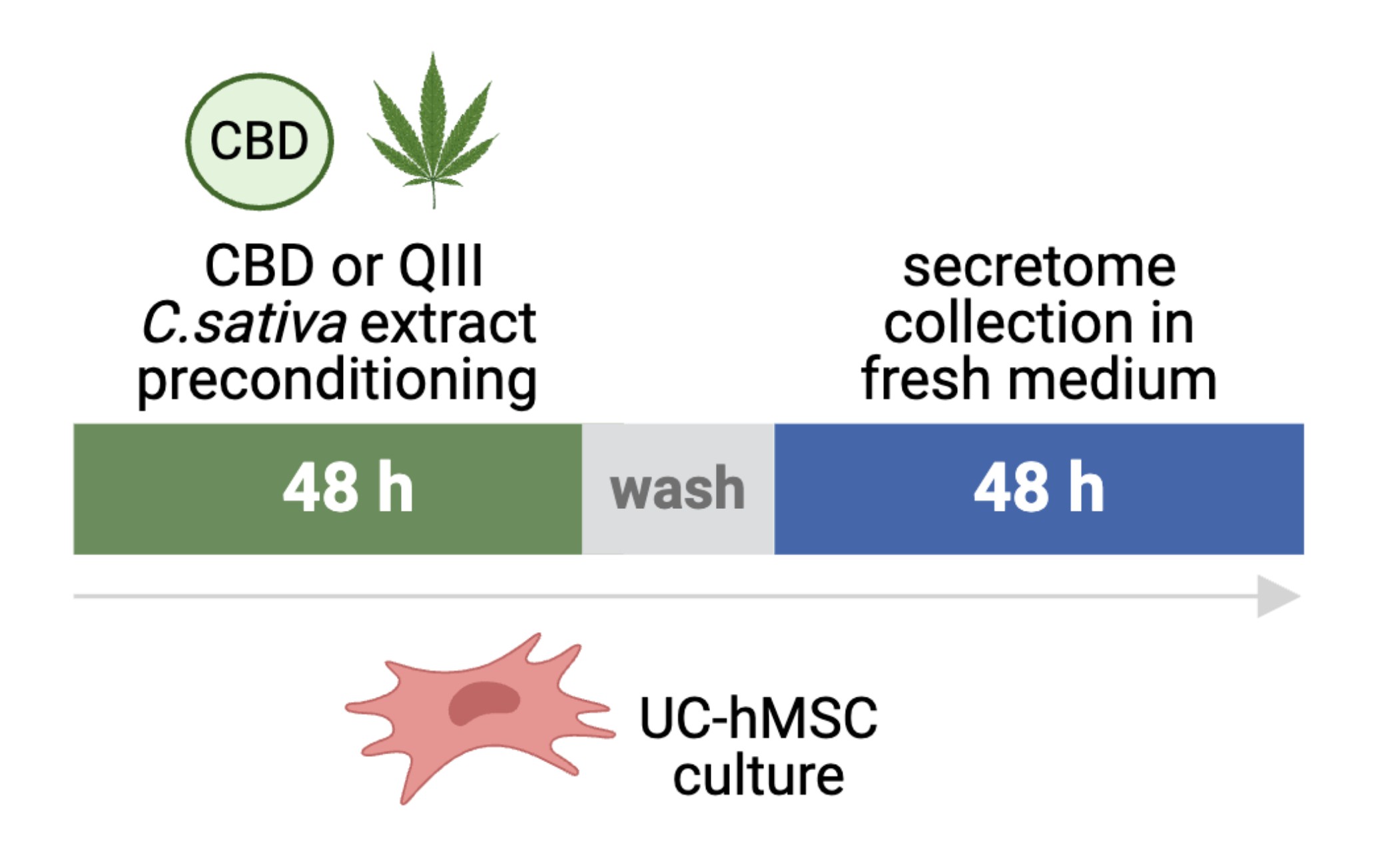

### Supplemental Figure 2

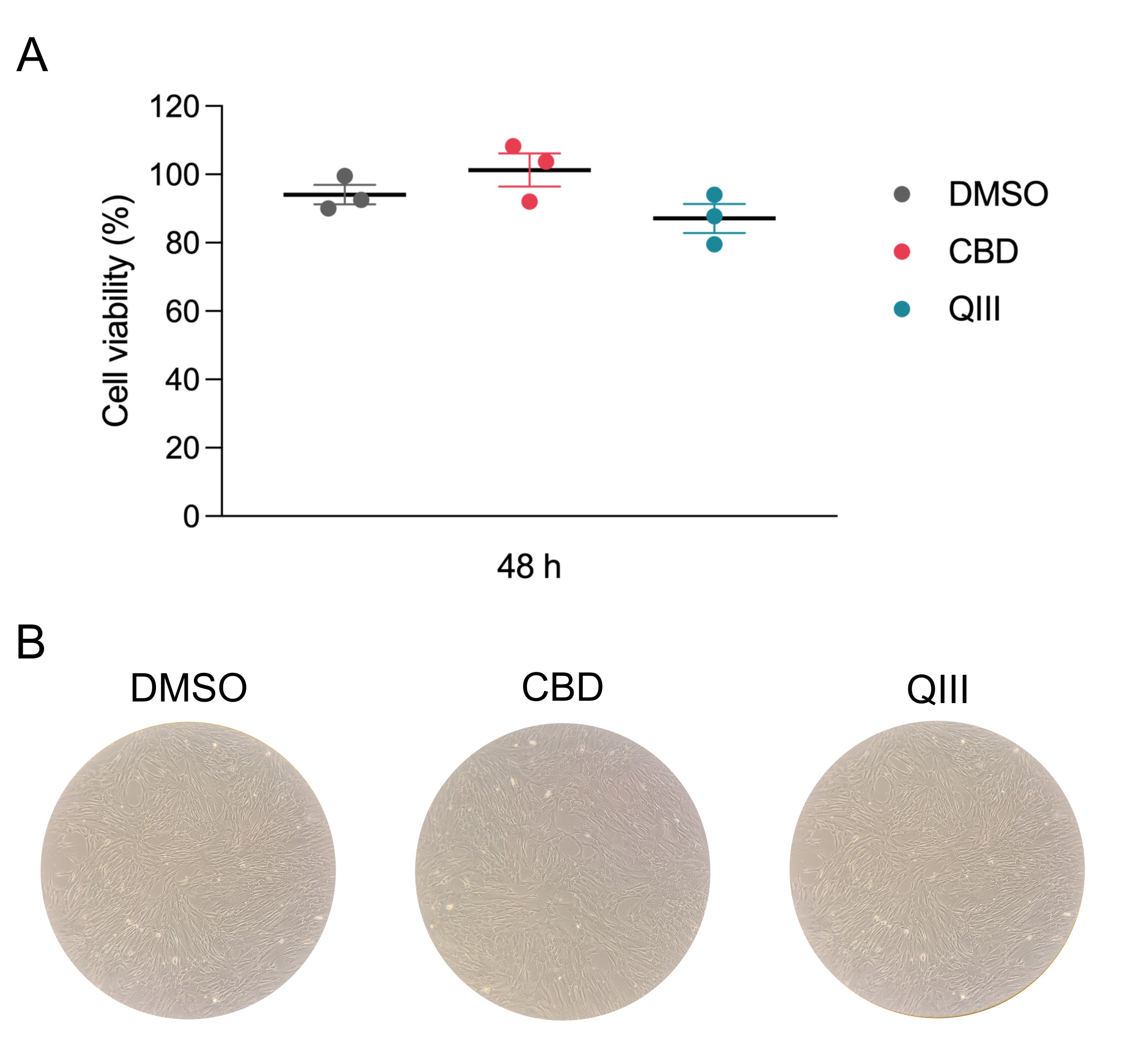

### Supplemental Figure 3

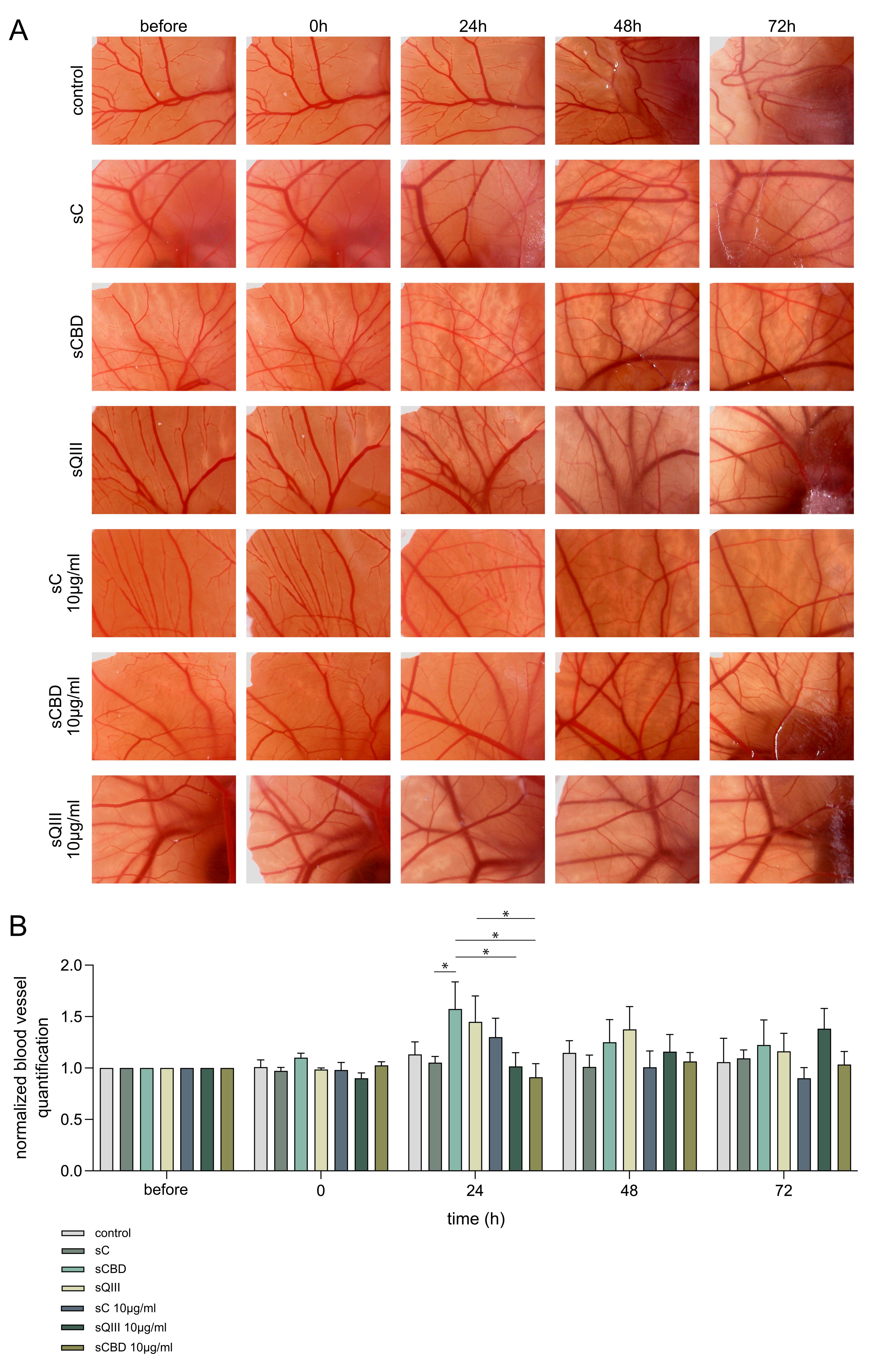
