## Supplementary Table 1 for "Purified CBD and CBD-rich full-spectrum *Cannabis sativa* extract potentiate the angiogenic paracrine function of umbilical cord derived mesenchymal stem cells"

The sequences of qRT-PCR primers (5′ to 3′)

| **Gene** | **Forward Primer** | **Reverse Primer** |
| --- | --- | --- |
| *PPARγ* | TCGAGGACACCGGAGAGG | GTGTCAACCATGGTCATTTCGTT |
| *TRPV-1* | AGAGTCACGCTGGCAACC | TGGAGGTGGCAGCATAATCG |
| *NAPE-PLD* | TTGTGAATCCGTGGCCAACATGG | TACTGCCATGGTGAAGCACG |
| *DAGLα* | AATGGCTATCATCTGGCTGAGC | TTCCGAGGGTGACATTCTTAGC |
| *DAGLβ* | GCGCAAAGTAAACGGCAAGA | CTGCAGCTTGGGCTTTTCAT |
| *VEGFA* | GCTCAGAGCGGAGAAAGCATT | ATTTACACGTCTGCGGATCTTG |
| *ANGPT1* | CTTGTGGCCCCTCCAATCTAA | GGCCCTTTGAAGTAGTGCCA |
| *EGF* | GGTGAGGAACAACCGCTACA | GGGAGCCTGAGCAGAAACTT |
| *NGF* | CTAAACAGCACACGGGGTGA | GAGCGCAGCGAGTTTTGGC |
| *CNTF* | CGCAGAGTCCAGGTTGATGT | CTGTAGCCGCTCTATCTGGC |
| *GDNF* | TGACTTGGGTCTGGGCTATGAAAC | TCGTACGTTGTCTCAGCTGCAT |
| *IDO* | TGGTAGCTCCTCAGGGAGAC | AAAGGCAACCCCCAGCTATC |
| *bFGF* | CCGGTCAAGGAAATACACCAGT | TATAGCTTTCTGCCCAGGTCCTGT |
| *SOD1* | GGTGGGCCAAAGGATGAAGAG | CCACAAGCCAAACGACTTCC |
| *SOD2* | GGAAGCCATCAAACGTGACTT | CCCGTTCCTTATTGAAACCAAGC |
| *GAPDH* | TTTTGCGTCGCCAGCCGAG | GACCAGGCGCCCAATACGA |
