## Supplementary Table 2 for "Purified CBD and CBD-rich full-spectrum *Cannabis sativa* extract potentiate the angiogenic paracrine function of umbilical cord derived mesenchymal stem cells"

Primary and secondary antibodies

| **Antibody** | **Species** | **Dilution** | **Company (Catalog#)** |
| --- | --- | --- | --- |
| phospho-ERK | rabbit | 1:1000 | Cell Signaling, #4370 |
| ERK | rabbit | 1:1000 | Cell Signaling, #4695 |
| VEGFA | rabbit | 1:1000 | Abcam, ab183100 |
| Phospho-VEGFR-2 | rabbit | 1:1000 | Cell Signaling, #2478 |
| α/β-tubulin | rabbit | 1:1000 | Cell Signaling, #2148 |
| HIF-1 α | rabbit | 1:1000 | Cell Signaling, #14179 |
| Anti-rabbit IgG | goat | 1:5000 | Jackson ImmunoResearch Laboratories, #111-035-003 |
